# eNODAL: an experimentally guided nutriomics data clustering method to unravel complex drug-diet interactions

**DOI:** 10.1101/2023.10.03.560662

**Authors:** Xiangnan Xu, Alistair M. Senior, David G. Le Couteur, Victoria C. Cogger, David Raubenheimer, David E. James, Benjamin Parker, Stephen J. Simpson, Samuel Muller, Jean Y.H. Yang

## Abstract

Unraveling the complex interplay between nutrients and drugs via their effects on ‘omics’ features could revolutionize our fundamental understanding of nutritional physiology, personalized nutrition and ultimately human health-span. Experimental studies in nutrition are starting to use large-scale ‘omics’ experiments to pick apart the effects of such interacting factors. However, the high dimensionality of the omics features, coupled with complex fully-factorial experimental designs together pose a challenge to the analysis. Current strategies for analyzing such types of data are based on between-feature correlations. However, these techniques risk overlooking important signals that arise from the experimental design and produce clusters that are hard to interpret. We present a novel approach for analyzing high-dimensional outcomes in nutriomics experiments, termed **e**xperiment-guided **N**utri**O**mics **D**at**A** c**L**ustering (**eNODAL**). This three-step hybrid framework takes advantage of both ANOVA-type analyses and unsupervised learning methods to extract maximum information from experimental nutriomics studies. First, eNODAL categorizes the omics features into interpretable groups based on the significance of response to the different experimental variables using an ANOVA-like test. Such groups may include the main effects of a nutritional intervention, and drug exposure, or their interaction. Second, consensus clustering is performed within each interpretable group to further identify subclusters of features with similar response profiles to these experimental factors. Third, eNODAL annotates these subclusters based on their experimental responses and biological pathways enriched within the subcluster. We validate eNODAL using data from a mouse experiment to test for the interaction effects of macronutrient intake and drugs that target aging mechanisms in mice.

## 1. Introduction

Nutrition is a powerful determinant of health and disease, but disentangling the single and interactive influences of nutrients and other dietary constituents poses considerable challenges, which are overlooked in conventional one-nutrient-at-a-time approaches (Simpson *et al*., 2015; Raubenheimer and Simpson, 2016). Adding to this complexity is the fact that nutritional requirements differ with genotype, development, infection and other environmental circumstances (Raubenheimer *et al*., 2022). Diet may also interact with non-nutritional factors such as drug treatments (Downer *et al*., 2020). Understanding how nutrients interact with one another and with such external factors to affect multiple levels of physiology and health is at the forefront of nutriomics, precision medicine and public health.

Pre-clinical nutrition science is now equipped with conceptual frameworks and multi-factorial experimental designs (Floyd *et al*., 2022), such as the geometric framework for nutrition (GFN) (Simpson *et al*., 2017), that can separate nutrient-nutrient and nutrient-non nutrient interactions and map response surfaces for different traits (from molecular to life-history responses) in *n*-dimensional nutrient space. Adding to this explanatory power is our ability to readily measure a myriad of ‘intermediary’ phenotypes as produced from large-scale ‘omics’ experiments. These outcomes generate insights into how experimental factors interact to determine health. The challenge now is how best to analyze the datasets produced from these multi-factorial experiments, where the number of omics features tends to be much larger than the sample size (Floyd *et al*., 2022; Thangadurai *et al*., 2022).

A common strategy to address this challenge is to group the high dimensional omics features into highly correlated clusters and then to analyze the relationship between these clusters and experimental factors. Examples of this approach are weighted correlation network analysis (WGCNA) (Langfelder and Horvath, 2008) and ClustOfVar (Chavent *et al*., 2012), which use unsupervised clustering of omics features based on correlation structure or their abundance value. Such methods have been widely used to analyze genomics and proteomics data (Pei *et al*., 2017). However, in the case of a multi-factorial nutritional experiment, these unsupervised clustering methods do not account for the experimental structure, therefore resulting clusters could be confounded with the study design. An illustrative example of this problem is as follows (shown in Figure S1). Consider the case where the abundance of two proteomic features, A and B, respond differently to the nutrient exposure in the presence of drug 1 *vs* drug 2. Despite responding differently to the experimental design, the marginal Spearman correlation of the two features can still be high (e.g., 0.74 in Figure S1). As a consequence, the majority of unsupervised learning algorithms would readily group these proteins together. A further complication of using unsupervised clustering methods in the context of experimental nutrition science, is that they do not provide biological interpretation of the resulting clusters, which makes it hard to understand how experimental factors affect the responses to feature clusters (Plant and Böhm, 2011).

We propose a novel statistical workflow for an **e**xperiment-guided **N**utri**O**mics **DA**ta c**L**ustering framework, which we coin eNODAL. This eNODAL workflow first uses an ANOVA-like model to distinguish whether an omics feature (e.g., a protein) shows significant response to the experimental design such as additive effects of a nutritional intervention (e.g., dietary carbohydrate) and some other external factor (e.g., drug exposure, genetic manipulation), or their interaction. Subsequently, a consensus clustering method is performed to further identify subclusters of features with similar response profiles. Finally, these subclusters are annotated based on both experimental response and pathway enrichment. This hybrid framework aims to capture both the effects of experimental treatments, and similarities in the profiles of molecular features. Using data from a recent multi-diet GFN study in mice (Le Couteur *et al*., 2021), we demonstrate how eNODAL clusters proteomics features based on their response to an experiment involving drug-diet interactions, and can then link these features to key phenotypes related to metabolic health. The data come from a complex experimental design consisting of ten different diets varying systematically in macronutrient ratio and energy density and three drug treatments with a control group. Using eNODAL we identify 29 interpretable proteomics subclusters representing different responses to nutrient intake, drug exposure and their interaction (i.e., proteins whose response to nutrient intake was substantially altered by drug exposure). Demonstrating the power of eNODAL, one such interactive subcluster, comprising proteins that are involved in the key AMPK pathway, would not have been identified via ANOVA or correlation-based clustering methods alone.

## 2. Material and Methods

### 2.1 Data

The data used come from an experimental study on the interactive effects of dietary macronutrients and gerotherapeutic drugs in mice (Le Couteur *et al*., 2021). In summary, male C57BL/6J mice were kept on one of ten different diets. The diets were designed to span across multidimensional nutrient space (protein, carbohydrate, fat, energy density), using the geometric framework for nutrition (GFN). Each diet comprised one of five different ratios of macronutrients (i.e., % energy from protein, carbohydrate, and fat), and were replicated at two energy densities (8 kJ/g and14.8kJ/g), with cellulose being used as the indigestible bulking agent to control energy density. Layered over this multidimensional nutritional design, animals were also on a control (no-drug) treatment or one of three gerotherapeutic: metformin, rapamycin, or resveratrol. In total, the fully factorial experimental design consisted of 40 different treatment combinations. Key metabolic traits, food intake and the intake of individual macronutrients were measured, and the abundance of the liver proteome was quantified.

The dataset has many of the challenges common to nutritional-omics datasets. First, there is a large number of outcomes; ca. 5,000 proteins were quantified. Second, the effects of multiple nutritional dimensions were captured; the experiment utilized 10 diets that covaried in protein, carbohydrates, fat content and overall energy density. Third, interactions between nutritional and non-nutritional factors were explored; each diet was replicated across three different drug treatments and one control. Finally, given the logistical constraints in performing dietary experiments the sample size precluded the application of complex machine-learning approaches.

### 2.2 Experiment-guided nutriomics data clustering (eNODAL) method

eNODAL hierarchically groups high-dimensional omics features guided by experimental factors (Figure 1 and Supplementary Figure S1). eNODAL has three key steps: an ANOVA-like test categorizes omics features into interpretable groups based on significant effects of treatments and/or their interactions (Section 2.3.1). Second, a consensus clustering method further divides these interpretable groups into subclusters to reflect distinct patterns of omics features (Section 2.3.2). Finally, these subclusters of features are annotated in two ways: (1) experimental responses (Section 2.4.2), and (2) pathway enrichment (Section 2.4.3).

**Figure 1:**
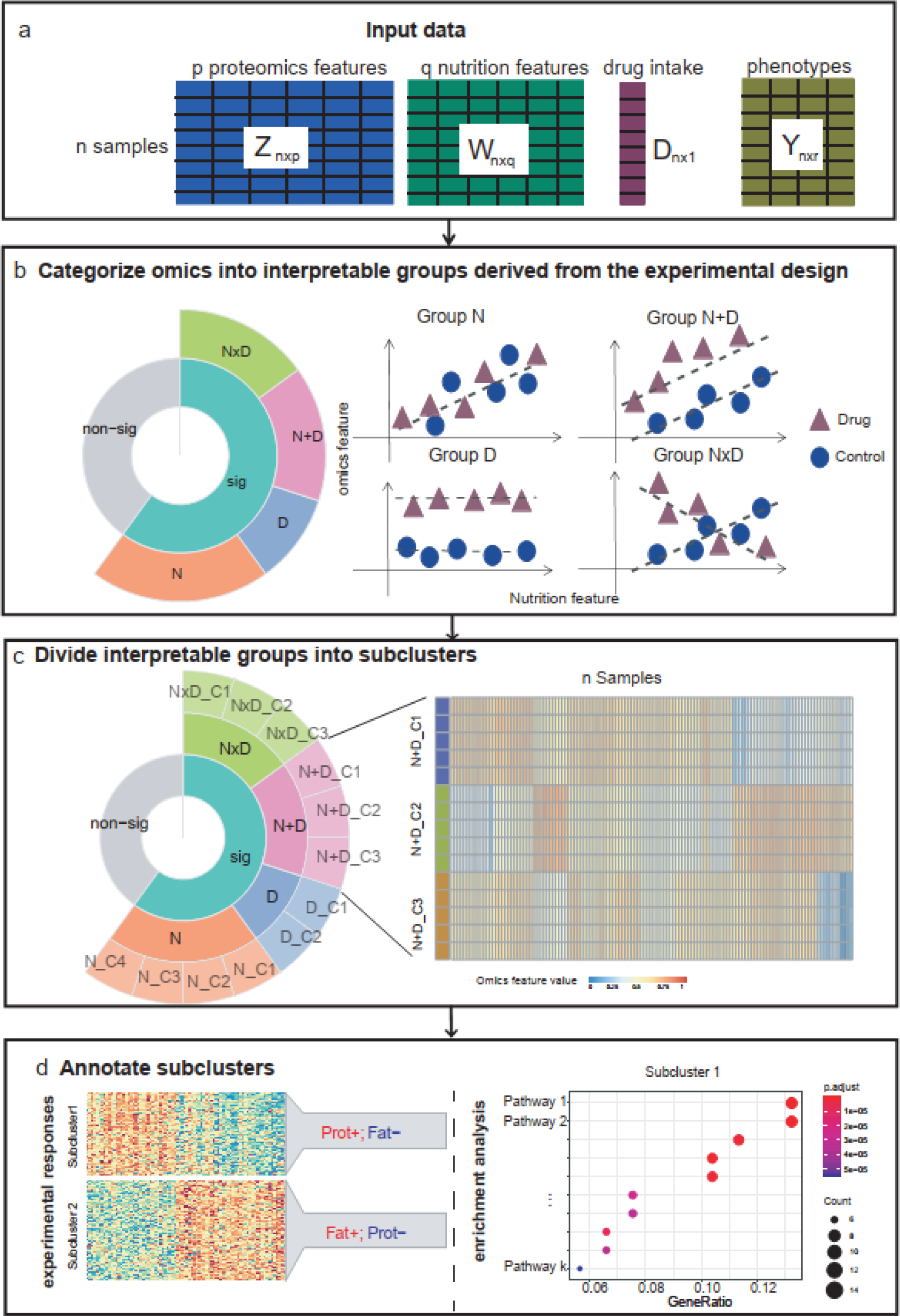
A schematic workflow for eNODAL showing four different stages. **a** Input of eNODAL including nutrition data, drug intake, omics data and metabolic phenotypes; **b** Categorize omics features into interpretable groups derived from the experimental design by ANOVA-like test; **c** Divide interpretable groups into subclusters via ensemble clustering; **d** Annotate subclusters based on their experimental responses and pathway enrichment analysis.

The first step is an ANOVA-like test that aims at categorizing the omics features to determine whether they show significant response to experimental factors as well as their interaction. This step classifies high-dimensional proteomics features into interpretable groups based on how they respond to nutrition, drug treatment and their interaction. Then a consensus clustering further divides these features into subclusters with similar profiles. Our consensus clustering is based on four types of distance measurement and four different clustering methods to provide robust results. After the first two steps, eNODAL provides two ways to annotate these subclusters from different perspectives: one angle is the relationship between nutrition intake and drug treatment, which is based on three sets of interpretable features derived from their response to experimental factors; another perspective is based on the biological pathway information, which is done by a KEGG pathway enrichment analysis. We will describe the details of each step in the following sections.

### 2.3 Two-stage clustering

The eNODAL framework uses a two-stage clustering method to group the high-dimensional omics features into subclusters. We will describe the details of each stage in the following sections.

#### 2.3.1 ANOVA-like test

The development of the first step ANOVA-like test is inspired by a nonparametric ANOVA (NANOVA) method which was first proposed to classify genes into different groups based on their factor effect (Zhou and Wong, 2011). We extend this method to categorize the proteomics features based on their response to a group of continuous variables (nutrition features) and a four-level categorical factor (drug treatment). We further consider the relationships among nutrition, treatment, and proteomics features. The non-linear version can be found in Supplementary Notes. We define the five nested models (M1, M2, …, M5) as follows:

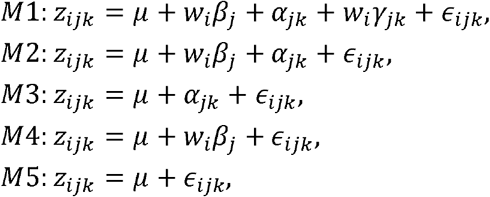

where *z*_*ijk*_ is the *j*^*th*^ proteomics feature for the ith sample which received the *k*^*th*^ gerotherapeutic drugs (*k* = 1 represents control group); *µ* is the overall effect; *α* _*jk*_ is the *k*^*th*^ treatment effect on *j*^*th*^ protein (in the mouse nutrition study, we have four different treatments corresponding to different drug intake); *w*_*i*_ is the nutrition features of *i*^*th*^ sample and *β* _*j*_ and *γ* _*jk*_ are the effect size of the relationships between nutrition and proteomics features. For *M*1, *β* _*j*_ and *γ*_*jk*_ are used to account for the main effects of treatment and nutrition as well as their interaction. For *M*2, *β*_*j*_ represent the contribution by nutrition intake. Proteins of *M*2 are affected by both nutrition and drugs, but their effects are independent.

Next, we categorize all proteins into five interpretable groups. Each group corresponds to one of the above five nested models, that is *M*1,*M*2,…,*M*5,, based on ANOVA-like testing as described below. We denote the set of all proteomics features as *S*.

The ANOVA-like testing proceeds as follows:

1. First, we identify proteins whose abundance are affected by either nutrition or treatment. This is achieved by the LC-test (Xu *et al*., 2020), which tests whether the effect of nutrition and treatment is significantly different from randomly permuted protein abundance. Proteomics features that show a significant response are assigned to the cluster “sig”, denoted as *C*_0_, and otherwise are assigned to the cluster “non-sig” (*S/C*_0_).
2. For the proteins in cluster “sig”, we use a nested ANOVA test to test whether the interaction effect in *M*1 is significant. That is, we test for each proteomics feature, H_0_: *γ*_*j*1_ = *γ* _*j*2_ = *γ*_*j*3_ = *γ*_*j*4_ = 0 versus *H*_1_ : at least one *γ* _*jk*_ is not equal to zero. The set of proteins with a significant interaction effect is denoted as *C*_*int*_ ⊂ *C*_0_.
3. For the proteins in set *C*_0_\ *C* _*int*_,we fit M2 and test whether coefficients *α*_*jk*_ (i.e. for each *j, H*_0_: *α*_*jk*_ = 0 = ∀k vs. *H*_1_ : at least one *α*_*jk*_ ≠ 0) and *β* _*j*_ (i.e. for each *j, H*_0_ : *β*_*j*_ = 0 vs. *H*_1_: *β*_*j*_ ≠ 0) is significant. Such a test also can be done via nested ANOVA tests in linear models. Proteins with *α*_*jk*_ ≠ 0 and *β* _*j*_ = 0 are classified as cluster “D”, denoted as *C*_D_, and those with *α*_*jk*_ = 0 and *β*_*j*_ ≠ 0 are classified as group “N”, denoted as *C*_*N*_ .
4. All proteins in *C*_0_\ (*C*_*int*_ ⋃ *C*_*N*_ ⋃ *C*_D_) form the “N+D” group.

After fitting the models and calculating the p-values, we use a hierarchical p-value adjustment to correct the p-value. Then the Bonferroni method is used to control the false discovery rate. Through this procedure, we classify the proteins into five interpretable groups, i.e. “NxD”, “N+D”, “N”, “D”, and “non-sig”.

#### 2.3.2 Consensus clustering

Based on the categorized five interpretable groups, we further divide the groups into subclusters using unsupervised clustering methods. We use a consensus clustering method with different types of distance measurements (Supplementary Notes) and varieties of clustering methods including affinity propagation (Frey and Dueck, 2007), Louvain clustering based on k-nearest neighbor graph (Blondel *et al*., 2008), a dynamic tree cut method for hierarchical clustering (Langfelder *et al*., 2008) and Density-based spatial clustering of applications with noise (DBSCAN) (Ester *et al*., 1996). These methods use a data-driven way to find the number of clusters and adapt well to the complexity of individual datasets.

Then a consensus matrix is created based on each individual clustering result. A binary similarity matrix is constructed from the corresponding clustering labels: if two features belong to the same cluster, their similarity is 1; otherwise, their similarity is 0. Finally, the resulting consensus matrix is clustered using the Louvain algorithm to get the resulting subclusters for each interpretable group.

### 2.4 Subcluster annotation

After two-stage clustering, we annotate these subclusters from two perspectives: first, from three sets of interpretable features described in Section 2.4.1; second, from pathway enrichment analysis. The details of the annotations of each subcluster are presented below.

#### 2.4.1 Calculate interpretable features for each protein

Let denote the *z*_*ijk*_ proteomics measurement in the *k*^*th*^ drug treatment group (*k* = 1,…,4, where *k* = 1 represents the control group), the corresponding *l*^*th*^ four nutrition nutrition intake is denoted *w*_*lk*_, and *z*_*j*_ = (*z*_*j*1_, *z*_*j*2_, *z*_*j*3_, *z*_*j*4_),w*l =* (*w*_*l*1_,*w*_*l*2_, *w*_*l*3_, *w*_*l*4_). In the mouse nutrition study, we focus on four nutrition intake features *l* =1,2,3,4, i.e. raw food intake in grams, and protein, carbohydrate, and fat intake in kJ. Three sets of interpretable features for the *j*^*th*^ proteomics measurements are described in the following.

**Set 1:** We first calculate the Fisher’s z-test statistic, *z*_*jl*_ (1,2,3,4) from the correlation coefficients of proteomics feature *z*_*j*_ and nutrition feature *w*_*l*_:

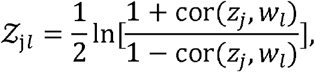

where cor (*Z*_*j*_,*w*_*l*_)is the sample correlation coefficient between protein *z*_*j*_ and nutrition *w*_*l*_. Each interpretable feature in Set 1 is calculated by 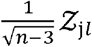 where *n* is the number of observations.

**Set 2:** We then calculate the pairwise t-statistic of differential abundance of *z*_*jk*_ between control (*k* = 1) and each treatment group (*k* = 2,3,4):

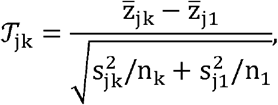

where 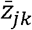 and 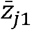 are the sample mean of protein abundance in drug group *k* and the control group, respectively; *S*_*jk*_, *S*_*j*1_ and *n*_*k*_, *n*_1_ are the corresponding sample standard deviation and sample size respectively. Set 2 is composed of *𝒯*_jk_, *k* = 2,3,4.

**Set 3**: Fisher’s z-test statistic of differential correlations of *z*_*jk*_ and *w*_*lk*_ between the control (*k* = 1) and treatment groups (*k* = 2,3,4) are calculated:

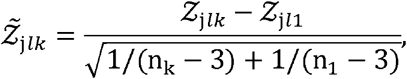

where 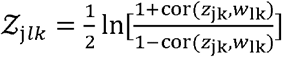 is the z-statistic similar to Set 1 but only uses a subset of the data in each drug group. Set 3 is a collection of 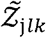, *l* = 1,2,3,4 and *k* = 2,3,4.

Set 1 shows the overall relationship between proteomics features and nutrition features.

Set 2 describes how liver protein abundance marginally changes with respect to different drugs.

Set 3 represents the change of the relationship between nutrition and proteomics in the three-drug treatment groups, i.e. metformin, rapamycin, or resveratrol, which showed to have an interaction effect between nutrition and drug.

#### 2.4.2 Annotate subclusters by interpretable features

The three created sets of interpretable features reflect different aspects of the relationship between proteomics features and experimental factors. We further annotate each subcluster based on these features.For the *J* subcluster, we take the annotation of its interaction effect, as an example: we first transform the related interpretable features 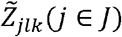 to ℱ_*jlk*_ as follows,

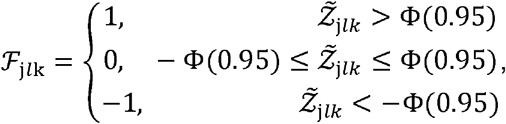

where Φ (·) is the cumulative distribution function of the standard normal distribution and Φ (0.95) ≈ 1.68. Then we calculate the proportion of *ℱ* _j*l*k_ = 1 or ℱ_j*l*k_ = −1 for proteins within subcluster *J*, i.e.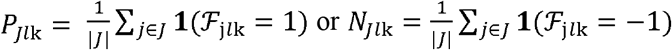, where **1**(·) is the indicator function. If *P*_*Jl*k_ > 0.7, it indicates that at least 70% of proteins in subcluster *J* show a significantly increased correlation with nutrition variable *l* in the drug group *k* compared with the correlation coefficients in the control group. Then we annotate cluster *J* with “increased correlation with variable in the drug group *k* “. A similar annotation procedure works for *N*_*Jkl*_ > 0.7.

#### 2.4.3 Annotate subclusters by pathway enrichment analysis

On the other hand, we also annotate each subcluster based on the enrichment of pathways in this cluster. This is done by enrichment analysis with the R package clusterProfiler (Wu *et al*., 2021) and the top enriched KEGG pathway is also used to describe each subcluster.

## 2.5 Creating network among proteins, subclusters and phenotypes

We first calculate the Spearman correlations between the *j*^*th*^ proteomics feature and the *m*^*th*^ metabolic phenotype, denoted as *ρ*_*jm*_ . The p-value for testing against *H*_0_: *ρ*_*jm*_ *=* 0 is calculated. If the p-value is smaller than 0.01, the corresponding Spearman correlation is set to zero. Then we use a gene set enrichment analysis (GSEA) like the multiset test method (Newton and Wang, 2015) to determine the significance of the correlation between a subcluster and selected metabolic phenotype. If the p-value is smaller than 0.01, we put an edge to emphasize the link between the corresponding subcluster and phenotype. Proteomics features and subclusters are linked by proteins showing high correlation (rank top 5) with the first principal component of proteins in the subclusters. The resulting network is drawn using the R-package ggnetwork (Tyner *et al*., 2017).

## 3. Results

With reference to our example mouse data, the experimental design yields five interpretable groups at the first eNODAL step. These groups have the following main and interaction effects arising from the experimental design: 1) “NxD” group: effects of a nutritional dimension and drugs, as well as their interaction; 2) “N+D” group: the additive nutritional and drug main effects; 3) “N” group: nutrient main effect only; 4) “D” group: drug main effect only; 5) “non-sig” group: intercept only group. Different types of visualization strategies used for these interpretable groups better suit their response to experimental factors (Figure S2). In the next step, a consensus clustering strategy (Kiselev *et al*., 2017) that integrates a variety of clustering methods and distance measurements is applied to the proteomics features in each of the 5 clusters in step 1 (Methods). Besides the widely used correlation distance (Kiselev *et al*., 2017; Langfelder and Horvath, 2008), two weighted correlation distances are proposed to account for the similarity in terms of the roles of nutrition intake, drug treatment, and their interactions (Supplementary Notes). Such weighted correlation distances dynamically focus on the correlations between variables as well as their contribution to experimental factors (Figure S3). Finally, we annotate these subclusters based on three sets of interpretable features (Methods) and enrichment analysis. In the mouse experiment, proteomics features are annotated with regard to response to macronutrient intake, drug treatment, and their interactions. The enrichment analysis provides functional biological annotation of these subclusters.

### 3.1 Categorizing omics data into interpretable groups derived from experiments

In the first step of eNODAL, we categorized the high dimensional proteomics features into interpretable groups based on whether they are significantly affected by diet, drug and/or interactions, with the results shown in Figure 2. A total of 2,951 proteins out of 4,987 proteins show significant responses to nutrient and/or drug exposure. Among these proteins with significant responses, the “N”, “N+D” and “NxD” groups are the majority groups with 1,350, 830, and 717 proteins, respectively, whereas the “D” group only has 53 proteins. The unbalanced number in each interpretable group implies nutrition shapes the largest fraction of the proteome. In contrast, a small number of proteins are affected solely by drug treatment (group “D”). That is not to say drugs have little effect on the proteome, rather those effects occur either additively, or in a more complex interaction, with diet (i.e., in groups “N+D” and “NxD”). Pathway analysis (Figure S4) shows that RNA splicing pathways are enriched in group “N” (rank 1, p<0.01) and “N+D” (rank 2, p<0.01), a finding consistent with our previous results (Le Couteur *et al*., 2021). For group “NxD”, the top enriched pathways are thermogenesis (p<0.01) and carbon metabolism (p<0.01). Several studies showed that thermogenesis is closely related to diet (Applied Research Press, 2015) and drug treatment (Peinado *et al*., 2014). Further, there is evidence suggesting that the interaction between drug and diet impacts thermogenesis (San-Cristobal *et al*., 2020). This implies that eNODAL can group proteomics features based on their response to experimental factors.

**Figure 2.**
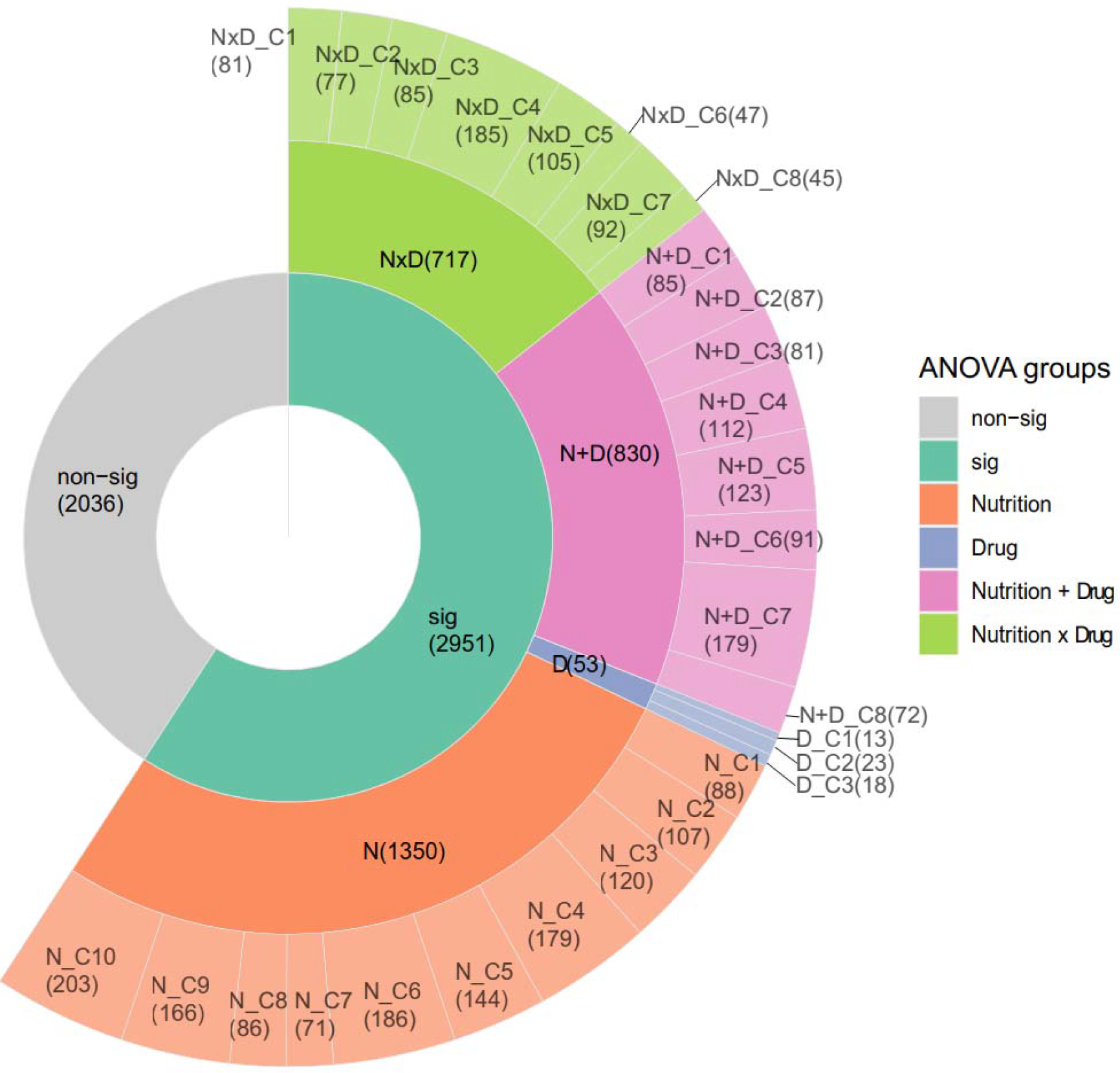
Clustering result of proteomics features from eNODAL. The 4,987 proteins are categorized into four interpretable groups based on an ANOVA-like test (two inner-layers). Then it is further clustered into 29 subclusters within each group (outer-layer). The numbers in each subcluster are shown within the round brackets.

### 3.2 Dividing interpretable groups into subclusters reveals different patterns of the proteomics features

To further identify clusters of proteins with similar patterns, the consensus clustering step of eNODAL subdivided the five broad groups of experimental responses. For the “N” group, we obtain nine subclusters. Figure 3 shows that these subclusters all have contrasting correlations with the different nutritional dimensions in the experiment (Figure 3a). For example, subcluster 5 in the “N” group (“N_C5”) comprises 144 proteins, the majority of which negatively correlate with total food intake in grams but positively correlate with carbohydrate and fat intake in kJ. Pathway analysis indicates that the Peroxisome pathway, which is known to be related to lipid metabolism (Le Couteur *et al*., 2021, Latruffe and Vamecq, 1997), is enriched in this subcluster of proteins. To visualize the effects of nutrient intake on within-subcluster protein abundance, we apply the surfaces-based approach from the GFN to the first principal component (PC1) of abundance within each cluster (Figure 3b-e). The subcluster “N_C5” (Figure S5) for example contains proteins with a higher abundance of elevated carbohydrate or protein intake, while the opposing pattern is seen in the subcluster “N_C6”. Similar results can be found within the much smaller “D” group, which is further clustered into three subclusters with different responses to drug treatment (Figure S6).

**Figure 3:**
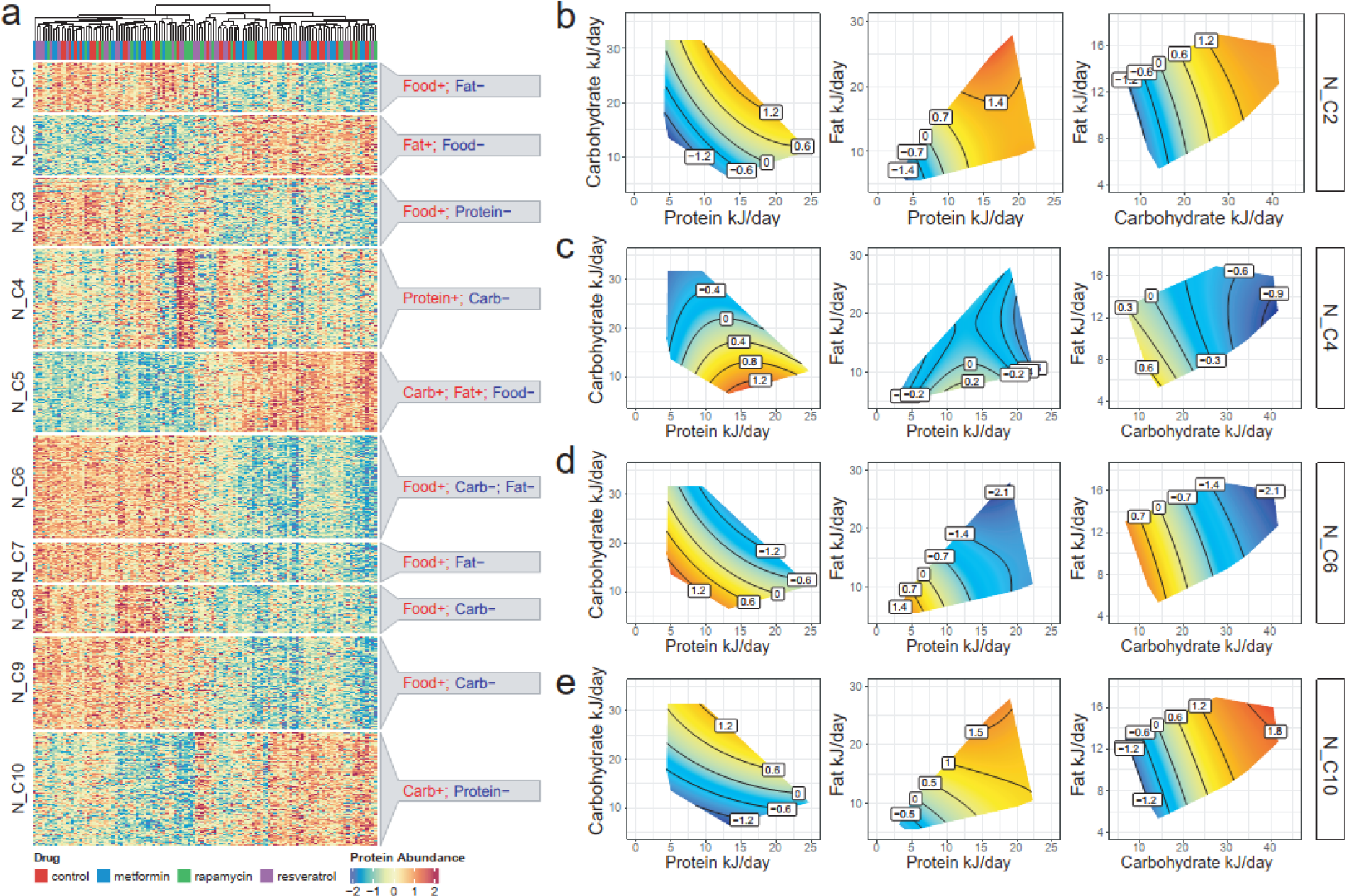
Main effect (N clusters): Subclusters in the “N” group and their annotations. **a** heatmap of the abundance of proteins in the “N” group, split by subclusters and annotation of each subcluster. **b-e** GFNs of the first PC of four subclusters in the “N” group. GFN of PC1 of subclusters “N_C2”, “N_C4”, “N_C6”, and “N_C10” respectively.

### 3.3 eNODAL reveals complex interplay among diet, drug and metabolic pathway

Both the “N+D” and the “NxD” group contain eight subclusters (Figure 4a and S7). In the “N+D” group, the effects of nutrition intake and drug treatment are “additive” (as denoted by the “+” sign). Here, a combination of GFN surfaces and boxplots can be used to visualize associations between nutrition intake as well as drug treatment and the proteomics features in the cluster (Figure S7).

**Figure 4:**
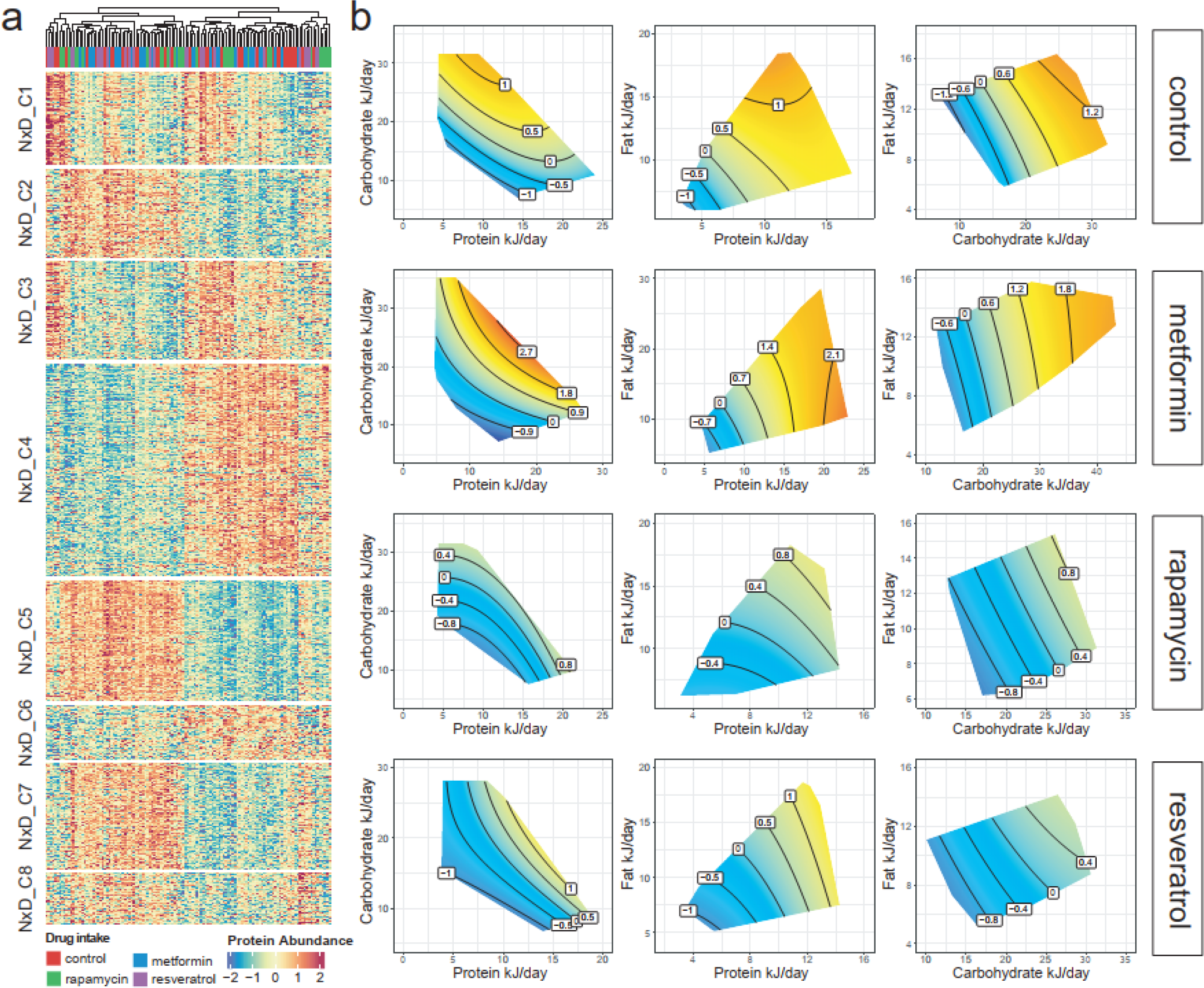
Interaction effect (“NxD” clusters): Subclusters in the “NxD” group and their annotations. **a** heatmap of the abundance of proteins in the “NxD” group, split by subclusters. **b** GFNs of the first PC of subcluster “NxD_C4”. GFNs are fitted based on the samples for each drug treatment respectively, resulting in 3 (combinations of nutrients intake) x 4 (number of treatments) =12 GFNs.

For the “NxD” group, the interaction effect between nutrition and drug contributes to the abundance of proteins in these subclusters (i.e., association between drug and protein abundance is dependent on nutrient intake). For proteins in the “NxD” groups, interpretation of the effects of the drug need to be evaluated with respect to the nutritional context. We visualize the association between nutrient intake and within cluster protein abundance (based on PC1 for the cluster) using the GFN surfaces visualized for each drug group separately (e.g., Figure 4b). Subcluster 4 in the “NxD” group (“NxD_C4”) contains the largest number of liver proteins (see Figure 4a). For “NxD_C4” proteins, increasing energy intake leads to elevated abundance of protein, but presence of rapamycin and resveratrol dampens this response. In the meantime, we also observed an effect related to protein-carbohydrate ratio (P:C) in the control group, say when holding energy constant, increasing P:C tends to reduce abundance of these proteins. While in drug groups, such P:C effect essentially disappears. We also see that the AMPK, insulin, and glucagon signaling pathways are enriched in this subcluster (see Figure S8). This result is consistent with previous studies where the activation of AMPK, a nutrient-sensing pathway, has been related to the intake of metformin (Wang *et al*., 2017) as well as interactions between diet and metformin (Marsh *et al*., 2010; Fu *et al*., 2015).

### 3.4 Network analysis reveals interplay among hub protein, subclusters and metabolic phenotypes

We jointly examined relationships between proteomics features, subclusters, and diet-related metabolic phenotypes. This step directly addresses our aim of understanding how diet by drug-affected proteins contributes to the metabolic phenotype and ultimately health of mice. This was achieved by creating a network to link proteomic features and metabolic traits and using multiset tests (Newton and Wang, 2015) to determine the significance of any identified associations (Methods). This analysis shows, for example, that the “NxD_C4” cluster shown in Figure 5 links closely with a large group of metabolic phenotypes that includes body weight, fasting insulin, and the mass of the retroperitoneal fat pad. Several other clusters of liver proteins that are positively affected by total energy intake also link to this cluster (e.g., “N_C2”, Figures 3b-e and 5). This result is consistent with previous findings (Le Couteur *et al*., 2021). A particular protein of note is Pex11, a hub protein in subcluster “NxD_C4”. Pex11 is positively correlated with many of the metabolic phenotypes (e.g., body weight and insulin levels) in our data and in previous studies (Chen *et al*., 2019). Examination of the GFN-type surfaces for this specific protein mirror those for “NxD_C4” as a whole (Figures 4b and S9), and Pex11 has been shown to be drug-diet responsive in other studies (Sharma *et al*., 2018; Li *et al*., 2002) .

**Figure 5:**
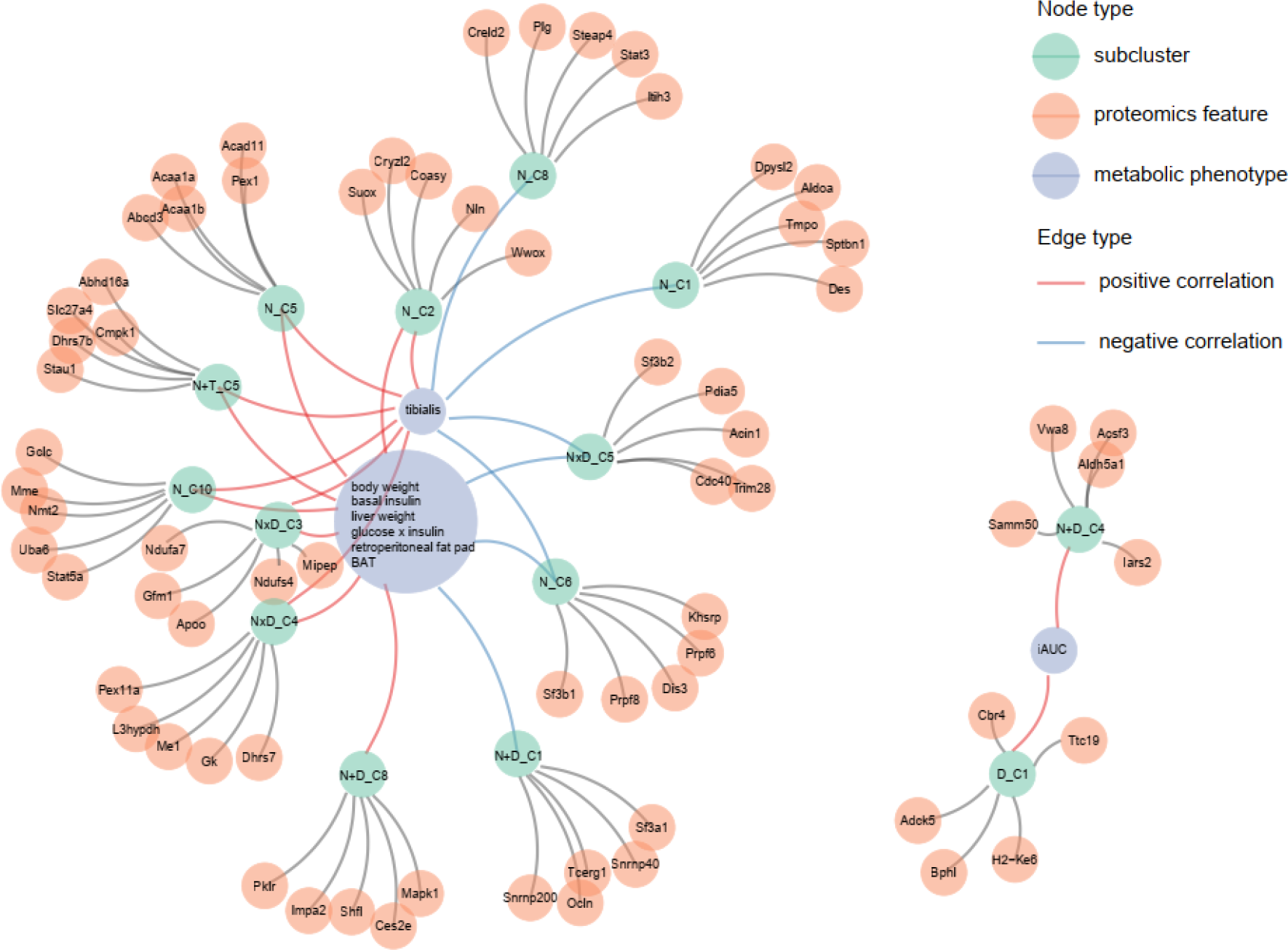
Network among hub proteins, subclusters and metabolic phenotypes: nodes represent subclusters, hub proteins and metabolic phenotype respectively. Edges between subclusters and proteins determined by top 5 proteins correlated with the first principal component of the subcluster. Edges between subclusters and metabolic phenotypes are determined by multiset test. Positive and negative correlation are calculated based on the correlation between median of the subcluster and metabolic phenotype.

We also observed that the incremental area under the curve for insulin (iAUC) is associated with two proteomics subclusters. Both subclusters contain proteins whose abundance is elevated under rapamycin exposure (“D_C1” and “N+D_C4”). Several previous studies have noted that rapamycin exposure decreases glucose tolerance in rodents (Yang *et al*., 2012; Weiss *et al*., 2018). Complementing this literature, eNODAL has identified that rapamycin may decrease glucose tolerance (i.e., increase the iAUC) by increasing the abundance of a suite of liver proteins (Figure 5).

## 4. Discussion

We present a three-step hybrid procedure called eNODAL, which integrates experimental structure with high-dimensional ‘omics’ features in fully factorial nutritional studies. This framework first categorizes the features into interpretable groups based on response to experimental treatments, before a consensus further divides these interpretable groups into subclusters with similar abundance profiles. Finally, we annotate these subclusters based on their experimental responses as well as enrichment of biological pathways. Demonstrating the power of eNODAL, we have analyzed data from a pre-clinical mouse experiment testing for interactions between diet and gerotherapeutic drugs affecting metabolic health and the liver proteome. Within these data, eNODAL obtained 29 subclusters of proteomics features representing different biological pathways. A number of these subclusters validate alternative analyses of the data, such as detecting the effects of the treatments on the spliceosome (Le Couteur *et al*., 2021). Furthermore, several of our results correlate with and complement findings from other studies on the effects of diet and gerotherapeutic drugs. For example, we see a negative effect of rapamycin on glucose homeostasis and demonstrate that these changes co-occur with effects of the drug on a cluster of specific live proteins.

When exploring an *n*-dimensional nutrition space, this flexibility is likely to be important. Several studies (Simpson and Raubenheimer, 2012; Xu *et al*., 2020) have detected associations between nutrient intake and gene expression that could be non-linear. We have therefore also implemented a hypothesis test using nonparametric generalized additive models (GAMs) (Wood, 2013, 2011; Wood *et al*., 2016; Solon-Biet *et al*., 2020; Senior *et al*., 2019) as well as a testing procedure to decide whether the use of a non-linear GAM significantly alters the results relative to using a linear model (see Supplementary notes). In our example dataset only 2% of proteins preferred GAM to the linear model. However, in other settings where many non-linear relationships exist, the use of GAM in the first stage is likely to be more appropriate. A further discussion about extension of eNODAL framework can be found in Supplementary notes.

The results from eNODAL provided more biological insights into the complex interplay between diet, drug, hepatic proteome and metabolic phenotype. On the one hand, eNODAL is able to identify RNA splicing pathways enriched in the “N” group, which were also found in our previous work. Furthermore, eNODAL identifies biological pathways related to interaction effects between nutrition and drugs, such as thermogenesis and AMPK pathways. Thermogenesis is closely related to the brown adipose tissue system and has shown its important role in regulation of body temperature (Kozak *et al*., 2010). Different types of diet, such as a high fat diet and/or high protein diet, as well as the intake of drugs may affect thermogenesis by altering metabolism (Applied Research Press, 2015; Clapham, 2012). AMPK pathway is also central to metabolic regulation, including energy production and storage and synthesis of fatty acids and cholesterol. The activation of AMPK pathways could be induced both by diet and drug intake (Agius *et al*., 2020; Song *et al*., 2018; Woods *et al*., 2017). Understanding the complex interplay among diet, drug as well as related metabolic pathways can help to optimize the effects of these substances on the regulation of the body system.

In summary, we present eNODAL, a three-step hybrid method to identify interpretable omics subclusters by integrating signals arising from complex nutritional experiments. The application of eNODAL on a complex mouse nutriomics study demonstrated its ability to identify biologically meaningful subclusters, such as subcluster that containing proteins enriched in AMPK signaling pathways, that are affected by interactions between nutrition and gerotherapeutic drugs, and not found (i.e. significantly enriched) by popular method WGCNA. eNODAL is implemented as an R package and can be downloaded freely from https://github.com/SydneyBioX/eNODAL.

## Availability and Implementation

eNODAL is implemented in R and the package is freely available at our github page: https://github.com/SydneyBioX/eNODAL

All nutrition data, phenotypical data, raw and processed proteomics data can be found in eNODAL R package (https://github.com/SydneyBioX/eNODAL) by command ‘data(“Proteomics_full”)*’*.

## Author contribution

JYHY, SM, AMS and XX conceived the study. XX led the method development and data analysis with input from SM and JYHY. ASM and XX led the evaluation of the method with input from all authors. DGLC, VCC, DR, DEJ, BP, and SJS provided the case study data and guided evaluation of the method. AMS analyzed and interpreted the nutriomics results. XX, AMS, JYHY and SM wrote the manuscript with input from all co-authors. All authors read and approved the final version of the manuscript.

## Acknowledgements

The authors thank all their colleagues, particularly at The University of Sydney, Sydney Precision Bioinformatics and Charles Perkins Centre for their support and intellectual engagement.

## Ethics approval

Ethical approval was granted by the Northern Sydney Local Health District Human Research Ethics Committee (HREC/18/HAWKE/109) and the North Shore Private Hospital ethics committee (NSPHEC 2018-LNR-009) and all participants provided written informed consent.

### Consent to participate

All participants provided written informed consent

### Consent for publication

All authors provide consent for publication

