## Supplementary information for "eNODAL: an experimentally guided nutriomics data clustering method to unravel complex drug-diet interactions"

**Supplementary Notes**

In the supplementary notes, we will first introduce the non-linear version of ANOVA-like test in the first stage. In addition, we will also introduce the test to determine whether to use linear model versus non-linear model. Then we describe the technical details of the consensus clustering procedure including the calculation of explained variance and calculation of weighted distance. Finally, we provide comparison between eNODAL result and widely used WGCNA clustering result for the mouse nutrition study.

**Non-linear ANOVA-like test**

Similar to the linear ANOVA-like test described in the manuscript, the non-linear version adopts the same procedure inspired from the nonparametric ANOVA (NANOVA) method (Zhou and Wong, 2011). We define the following five nested models:

$$M1': z_{ijk}=\mu+f_{j}(w_{i})+\alpha_{jk}+g_{jk}(w_{i})+\epsilon_{ijk},$$

$$M2':z_{ijk}=\mu+f_{j}(w_{i})+\alpha_{jk}+\epsilon_{ijk},$$

$$M3: z_{ijk}=\mu+\alpha_{jk}+\epsilon_{ijk},$$

$$M4':z_{ijk}=\mu+f_{j}(w_{i})+\epsilon_{ijk},$$

$$M5: z_{ijk}=\mu+\epsilon_{ijk},$$

All of the symbols have similar meaning as its linear counterpart, except we replace the linear terms $w_{i}\beta_{j}$ and $w_{i}\gamma_{jk}$ with non-linear functions $f_{j}(w_{i})$ and $g_{jk}(w_{i})$. As is in the usual ANOVA model, constraints $\sum_{k=1}^{4} \alpha_{jk}=0$ and $\sum_{k=1}^{4} g_{jk}(w_{i})=0\forall j$, are imposed for identifiability (Zhou and Wong, 2011).

Then, we also categorize all proteins into five interpretable groups. The procedure is the same as the linear version, we only replace the nested ANOVA test to non-linear ANOVA-like test. Specifically, we replace (2) and (3) in the manuscript with (2)’ and (3)’ as follows:

(2)’ For the proteins in cluster "sig", we use a nested ANOVA test to test whether the interaction effect in M1’ is significant. That is, we test for each proteomics feature, ${H_{0}:g}_{j1}=g_{j2}=g_{j3}=g_{j4}=0$ versus $H_{1}$: at least one $g_{jk}$ is not equal to zero. The set of proteins with a significant interaction effect is denoted as $C_{\mathrm{int}}\subset C_{0}$.

(3)’ For the proteins in set $C_{0}\backslash C_{int}$, we fit M2’ and test whether coefficients $\alpha_{jk}$ (i.e. for each $j$, $H_{0}:\alpha_{jk}$ = 0, k vs. $H_{1}$: at least one $\alpha_{jk}$ ≠0) and $f_{j}$ (i.e. for each $j$, $H_{0}$: $f_{j}$ = 0 vs. $H_{1}$: $f_{j}$ ≠ 0) is significant. Such a test also can be done via nested ANOVA tests in linear models. Proteins with $\alpha_{jk}$ ≠ 0 and $f_{j}$ = 0 are classified as cluster "D", denoted as CT, and those with $\alpha_{jk}$ = 0 and $f_{j}$ ≠ 0 are classified as group "N", denoted as $C_{N}$.

All other procedures keep the same as it is in the ANOVA-like test.

**Testing between linear model and non-linear model**

The testing procedure between linear model and non-linear model is also inspired from the NANOVA method (Zhou and Wong, 2011). We define the following two models:

$$M6:z_{ijk}=\mu+w_{i}\beta_{j}+f_{j}(w_{i})+\alpha_{jk}+w_{i}\gamma_{jk}+g_{jk}(w_{i})+\epsilon_{ijk},$$

$$M7:z_{ijk}=\mu+w_{i}\beta_{j}+\alpha_{jk}+w_{i}\gamma_{jk}+\epsilon_{ijk},$$

where the symbols have the same meaning as it is in the previous section. The null hypothesis is $H_{0}$: $g_{j1}=g_{j2}=g_{j3}=g_{j4}=f_{j}=0$ *vs* at least one of these functions not equal to 0. The testing is based on a bootstrapping procedure and describe as follows:

(1) Fit M6 and M7. The corresponding residuals for sample $i$ proteomics feature $j$ and treatment $k$ are denoted as ${\epsilon_{ijk}}^{(M6)}$ and ${\epsilon_{ijk}}^{(M7)}$ respectively.

(2) Calculate the F-like statistics for proteomics feature $j (j=1,...,p)$ by $F_{j}=\frac{\sum_{i=1}^{n} \sum_{k=1}^{4} [{(\epsilon_{ijk}}^{(M7)})^{2}-{(\epsilon_{ijk}}^{(M6)})^{2}]}{\sum_{i=1}^{n} \sum_{k=1}^{4} {(\epsilon_{ijk}}^{(M6)})^{2}}$

(3) Sampling ${\tilde{\epsilon}_{ijk}}^{(M6)} (i=1,...,n; j=1,...,p;k=1,...,4)$ with replacement from ${\epsilon_{ijk}}^{(M6)}$.

(4) Let ${z^{*}}_{ijk}=z_{ijk}- {\epsilon_{ijk}}^{(M7)}+{\tilde{\epsilon}_{ijk}}^{(M6)}$. Fit M6 and M7 using ${z^{*}}_{ijk}$ and calculate the corresponding residuals ${\epsilon_{ijk}}^{*(M6)}$ and ${\epsilon_{ijk}}^{*(M7)}$ respectively.

(5) Calculate the F-like statistics for proteomics feature $j(j=1,...,p)$ by

${F_{j}}^{(1)*}=\frac{\sum_{i=1}^{n} \sum_{k=1}^{4} [{{(\epsilon}_{ijk}}^{*(M7)})^{2}-({\epsilon_{ijk}}^{*(M6)})^{2}]}{\sum_{i=1}^{n} \sum_{k=1}^{4} ({\epsilon_{ijk}}^{*(M6)})^{2}}$.

(6) Repeat (3)-(5) B times and get ${F_{j}}^{(2)*}$,...,${F_{j}}^{(B)*}$.

(7) Calculate the p-value for proteomics feature $j(j=1,...,p)$ by $p_{j}=\frac{1}{B}\sum_{b=1}^{B} \boldsymbol{1}( {F_{j}}^{(b)*}>F_{j}).$

Similar to the procedure in NANOVA, ${\tilde{\epsilon}_{ijk}}^{(M6)}$ is used to simulate the random error distribution and ${z^{*}}_{ijk}$ is generated from the null model by adding the resampled residuals to the estimated null model $z_{ijk}=\mu+w_{i}\hat{\beta}_{j}+\hat{\alpha}_{jk}+w_{i}\hat{\gamma}_{jk}$, where the $\hat{\beta}_{j}$, $\hat{\alpha}_{jk}$ and $\hat{\gamma}_{jk}$ are the estimated $\beta_{j}$, $\alpha_{jk}$ and $\gamma_{jk}$ respectively.

Instead of the standard F statistics as it is in NANOVA, we use the F-like statistics which are not corrected by the degree of freedom. In our case, since the degree of freedom is a constant in under observed $F_{j}$ and simulated ${F_{j}}^{(2)*}$, which do not affect the result of the empirical p-value calculated in step (7).

**Contribution of nutrition intake, drug treatment and their interaction**

To better describe the interpretable groups "N+D" and "NxD", we measure the contribution of groups of variables based on the difference of the residual sum of squares (RSS) between the full model and the reduced model, i.e. setting part of the parameters to zero. For group "NxD", we fit the following GAM in Equation (M1) - (M4). Then we calculate the difference of RSS between the full model and the reduced models:

$$\begin{matrix} {\Delta\mathrm{RSS}}_{\mathrm{int}}=RSS(M2)-RSS(M1), \\ {\Delta\mathrm{RSS}}_{N}=RSS(M3)-RSS(M2), \\ {\Delta\mathrm{RSS}}_{T}=RSS(M4)-RSS(M2). \end{matrix}$$

where $\mathrm{RSS}(M)$ is the $\mathrm{RSS}$ of model M. The contribution of the interaction term $V_{\mathrm{int}}$ is defined as the normalized $\Delta\mathrm{RSS}$, i.e.

$V_{int}=\frac{\Delta RSS_{\mathrm{int}}}{\Delta RSS_{\mathrm{int}}+\Delta RSS_{N}+\Delta RSS_{D}}$

A Similar calculation can be applied for the contribution of nutrition $V_{N}$ and drug intake $V_{D}$. We also apply it to the "N+D" group and get the contribution of nutrition features and drug intake for each proteomics feature.

**Distance for consensus clustering**

We use four different types of distance for our consensus clustering. The first two are the widely used Pearson and Spearman correlation distance (Kiselev *et al.*, 2017), which we denote as $D_{pcc}$and $D_{spc}$, respectively. We also use two types of weighted correlation distance to account for their similarity in terms of contribution of nutrition intake, drug treatment, and their interactions. We use $D_{pcc}$ as an illustration and the same calculation can be applied to $D_{spc}$. First, we calculate the Aitchison distance (Greenacre, M. 2021) of $(V_{N} ,V_{D} ,V_{int})$ as done in the previous section between pairs of proteomics features. We denote this distance matrix as $D_{V}$. Then we normalise^13^ $D_{V}$and $D_{pcc}$ by $\tilde{D}_{V}=\frac{D_{V}}{max(D_{V})}$ and $\tilde{D}_{pcc}=\frac{D_{pcc}}{max(D_{pcc})}$. The weighted Pearson distance is calculated by $D_{wpcc}=(1-\tilde{D}_{V})\tilde{D}_{pcc}+{\tilde{D}_{V}}^{2}$.

$D_{wpcc}$dynamically alternates the focus between $D_{V}$and $D_{pcc}$. As is illustrated in Figure S3, two pairs of points with different contributions of nutrition ("N"), drug treatment ("D"), and their interaction ("NxD") are shown. We observe that when the $(j_{1},j_{2})$th element in $\tilde{D}_{V}$ is close to 0 (two green points), then the roles of the experimental factors are similar for protein $j_{1}$ and protein $j_{2}$; additionally, $D_{wpcc}(j_{1},j_{2})$ is close to $\tilde{D}_{pcc}(j_{1},j_{2})$, as $D_{wpcc}(j_{1},j_{2})$ focuses more on the correlation structure. When $\tilde{D}_{V}$ is close to 1 (two pink points), which means that the contribution of some experimental factors shows the discrepancy between two proteins, then $D_{wpcc}$ tends to focus more on $\tilde{D}_{V}$.

**Visualization using Geometric Framework for Nutrition (GFN)**

To visualize the relationships between nutrition and proteomics features, we use a surfaces-based approach, geometric framework for nutrition (GFN), to illustrate such association accounts for their possible nonlinear interactions (Simpson *et al.*, 2017). GFN uses a general additive model (GAM) where omics feature as the response and nutrition features as the covariate. After fitting the model, the fitted function is used to generate response surfaces based on selected nutrition features.

Here, GFN visualization of each proteomics feature varies across the interpretable groups. For the "N" group, we use all samples to fit a GAM model based on $M4$ in Equation $M4$; the fitted value is used to visualize the response surface. For the "N+D" group, the GFNs are based on Equation $M2$ by subtracting the treatment effect in the model. For the "NxD" group, the fitted value of the response surface of the GFNs is based on Equation $M1$ within each drug treatment.

**Comparison between eNODAL and WGCNA**

We evaluate the clustering result between eNODAL and the result from WGCNA (Langfelder and Horvath, 2008), , a widely used method for categorizing high-dimensional features. In this mouse nutriomics dataset, WGCNA obtains 5 clusters of proteomics features, containing 1671, 970, 140, 97, and 73 proteins respectively.

The adjusted rand index (ARI), a cluster comparison statistic, is quite low (≈0.08) between clustering results from WGCNA and eNODAL. We use the subcluster annotation procedure (Methods) to annotate WGCNA clusters. It turns out most of the subclusters are driven by a marginal correlation between nutrition intake and proteomics abundance. As is shown in the previous result, macronutrients have a much larger effect on proteins than treatment. eNODAL, however, further reveals relationships among protein abundance, nutrition intake, and drug treatment in detail compared with WGCNA. Taking the WGCNA cluster 2 ("W_C2") as an example, the majority of its proteins show a negative correlation with raw food intake and a positive correlation with fat intake; while, in the eNODAL result, the majority of these proteins are distributed in subclusters "N_C2", "N_C5", "N_C10", "N+D_C4, "N+D_C5", "N+D_C8", "NxD_C4" and "NxD_C6" (Figure S10a). We noticed that proteins in subcluster "N+D_C5" not only share a similar marginal correlation with nutrition intake to "W_C2", but they also exhibit lower abundance in the resveratrol group. Similarly, subcluster "NxD_C4" further reveals the interaction between nutrient intake and drug treatment.

We notice that the insulin signaling pathway is enriched in "W_C2" (p<0.01, rank 3, Figure S10b). At the same time, this pathway is also enriched in eNODAL subclusters "N+D_C8"(p < 0.01, rank 1, Figure S10c) and "NxD_C4"(p < 0.01, rank 2, Figure S8). Except for the relationship with nutrient intake, proteins in subcluster "N+D_C8" show lower abundance in the rapamycin group and resveratrol group in "NxD_C4". Several studies pointed out that this pathway is affected by high carbohydrate or fat diet intake (Liu *et al.*, 2015; Li *et al.*, 2021) as well as rapamycin (Li *et al.*, 2010) and resveratrol (André *et al.*, 2017; Sadi *et al.*, 2015). It further indicates that subclusters by eNODAL can integrate the information from nutrition intake, drug treatment, and protein abundance and provide a biologically meaningful interpretation of these subclusters.

**Further discussion**

The first component of eNODAL incorporates experimental design information (e.g., factorial design structure) in the overall classification of omics features. In our demonstration analysis, we use an ANOVA-like F-test to determine the association between experimental factors and the abundance of each proteomics feature. This choice was motivated by the simplicity of the F-test. However, the package we have developed also implements other tests for linear models including global tests (Goeman *et al.*, 2004, 2006) and the Tmax test (Goeman *et al.*, 2006). For our case-study, these other options show similar results, but we highlight here that the eNODAL framework can provide different choices of hypothesis testing procedures to suit different data types.

The second component of eNODAL uses data-driven clustering to generate refined subclusters. These clustering methods are also able to be user-defined to accommodate data structure or prior biological knowledge. Our current strategy uses consensus clustering algorithms with varying distance measures. This multimodal approach ensures that the number of estimated clusters is stable and robust to differences between distance measures. In practice, if the number of subclusters is known through external biological knowledge, such information can be incorporated by using a form of unsupervised clustering such as *k-*means or hierarchical clustering.

The proposed eNODAL method was initially motivated by analyses of the effects of nutrition and drugs on the proteome. However, our approach is not restricted to the examination of these three data modalities. A natural extension of eNODAL would be to handle experiments generating multi-omics data. eNODAL can be easily adapted by extending both stages of eNODAL to handle multi-omics data. In the first component, we classify each omics feature according to whether it responds significantly to each experimental factor or their interactions. Then various multi-omics clustering algorithms, as described in a recent review (Rappoport and Shamir, 2019), can be used to modify the subclustering in the second component. For example, we can readily construct a similarity network for each omics feature inside each interpretable group before integrating and clustering multi-omics features using the network fusion (Wang *et al.*, 2014).

One of the key findings from our application of eNODAL to these data is the relatively small number of proteins that are affected by gerotherapeutic independent of the dietary context. In this study just 1.8% of significant proteins (i.e., 53/2,951) were affected by drugs alone, with no effect of diet. In contrast, the dietary treatment had many more effects, being responsible alone for 45.7% of significant effects. Importantly, around a quarter of the affected proteins demonstrated an interactive response between drugs and diet. Furthermore, these components of the proteome can be directly linked to metabolic phenotype. This result indicates that the effects of gerotherapeutic drugs on metabolic health are likely to be diet-dependent, and would possibly have heterogeneous effects if consumed by a wide population, due to dietary differences. Additional influences, e.g., genetics, will complicate matters further. On the basis of this result, we would caution against the widespread public uptake of such ‘health-span extending’ drugs (Lee *et al.*, 2021), at least until their diet synergies have been better mapped.

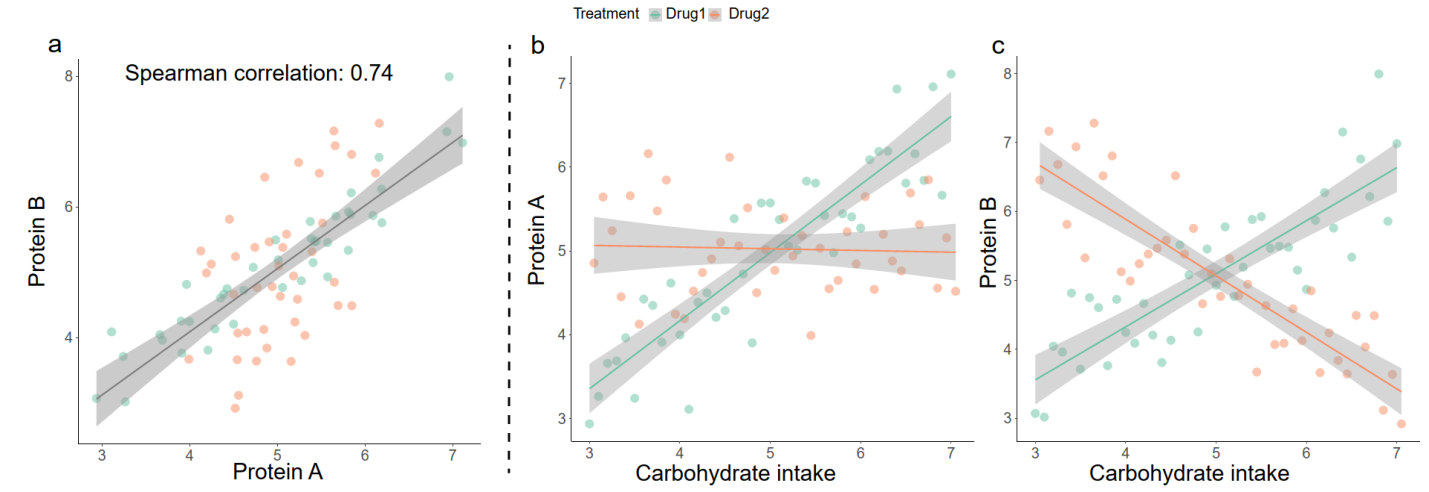

Fig S1. **A toy example illustrates that proteins with different responses to nutrition and drug intake could be highly correlated**. **a** Protein A and B are highly correlated. **b** Protein A showed a positive correlation with Carbohydrate intake in the presence of drug 1. **c** Protein B showed a different correlation with Carbohydrate intake with drug 1 and 2.

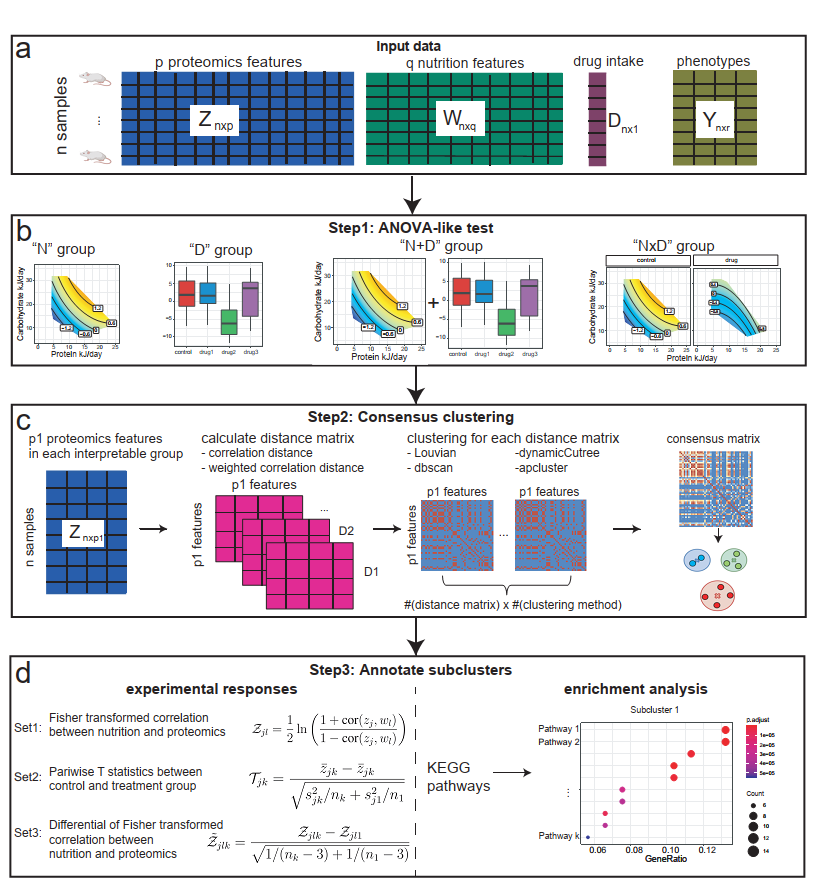
 Fig S2. **overview of algorithms in the eNODAL framework.** **a** Input data of eNODAL framework. **b** eNODAL first uses an ANOVA-like test to determine how each protein responds to experimental conditions. In our data, we have four groups, "N", "D", "N+D" and "NxD", which are significantly affected by nutrition, drug treatment, additive effect of nutrition and drug treatment and interaction effect of nutrition and drug treatment. Different visualization strategies are used for each group. We use GFN, boxplot, GFN and boxplot, and GFNs for each treatment group to visualize the effect of experimental factors on proteins for "N", "D", "N+D" and "NxD" groups respectively. **c** eNODAL uses a consensus clustering to further divide the interpretable groups into subclusters. It first calculates four different types of distance (pcc, spc, wpcc, wspc) between pairs of proteins and then four different clustering methods (Louvian, apcluster, dbscan and dynamicCuttree) derived a consensus matrix. Then a Louvian clustering on the consensus matrix used for the subclusters. **d** Annotation of these subclusters based on experimental responses and pathway enrichment analysis.

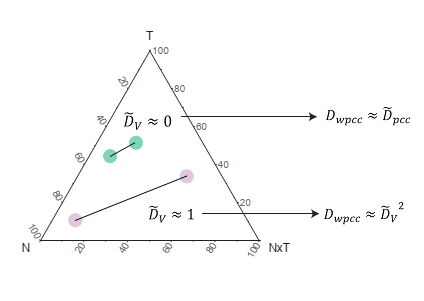

Fig S3. **Illustration of weighted Pearson distance.** The axes in the triangle show the contribution of nutrition ("N"), drug treatment ("D") and their interactions ("NxD").

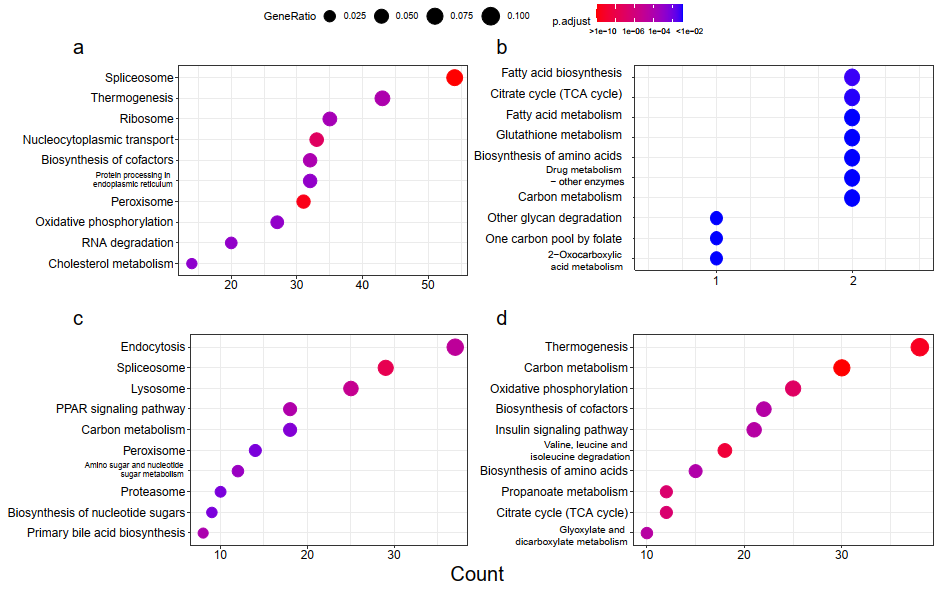

Fig S4. **KEGG pathway enrichment analysis of groups by ANOVA-like tests. a-d** KEGG pathway enrichment of the "N", "D", "N+D", "NxD" group respectively: y-axis represents the top 10 pathways with smallest adjusted p-value, x-axis represents the number of proteins in the corresponding pathway.

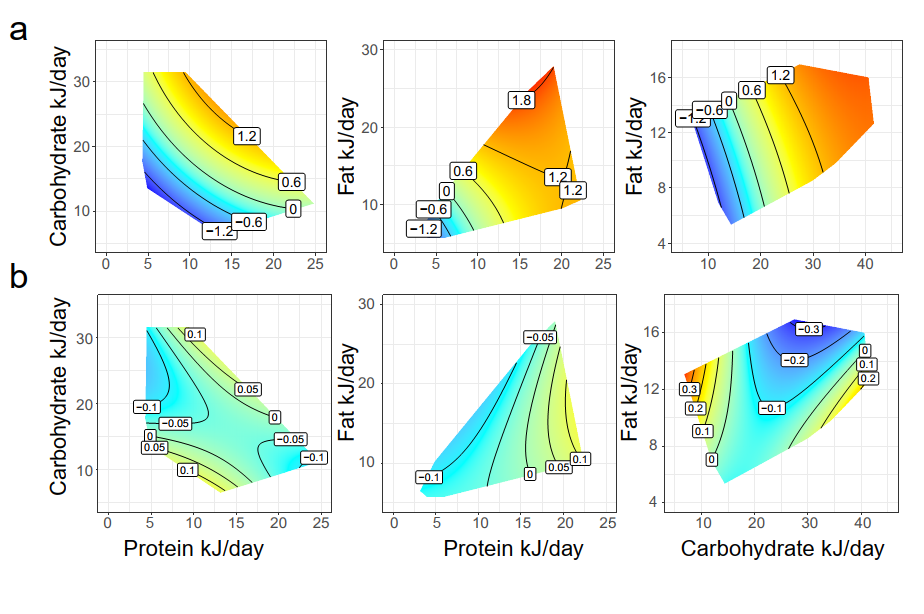

Fig S5. **GFNs of PC1 and PC2 for subcluster** "**N_C5**"**.** **a-b,** GFN of PC1 and PC2 for subcluster "N_C5" respectively.

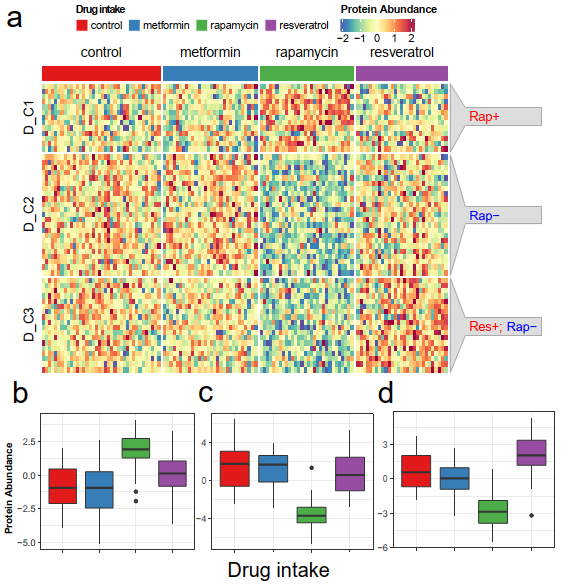

Fig S6. **Subclusters in the "D" group and their annotations.** **a** Protein abundance of three subclusters in "D" group, **b-d** boxplot of PC1 of subclusters "D_C1", "D_C2" and "D_C3" respectively.

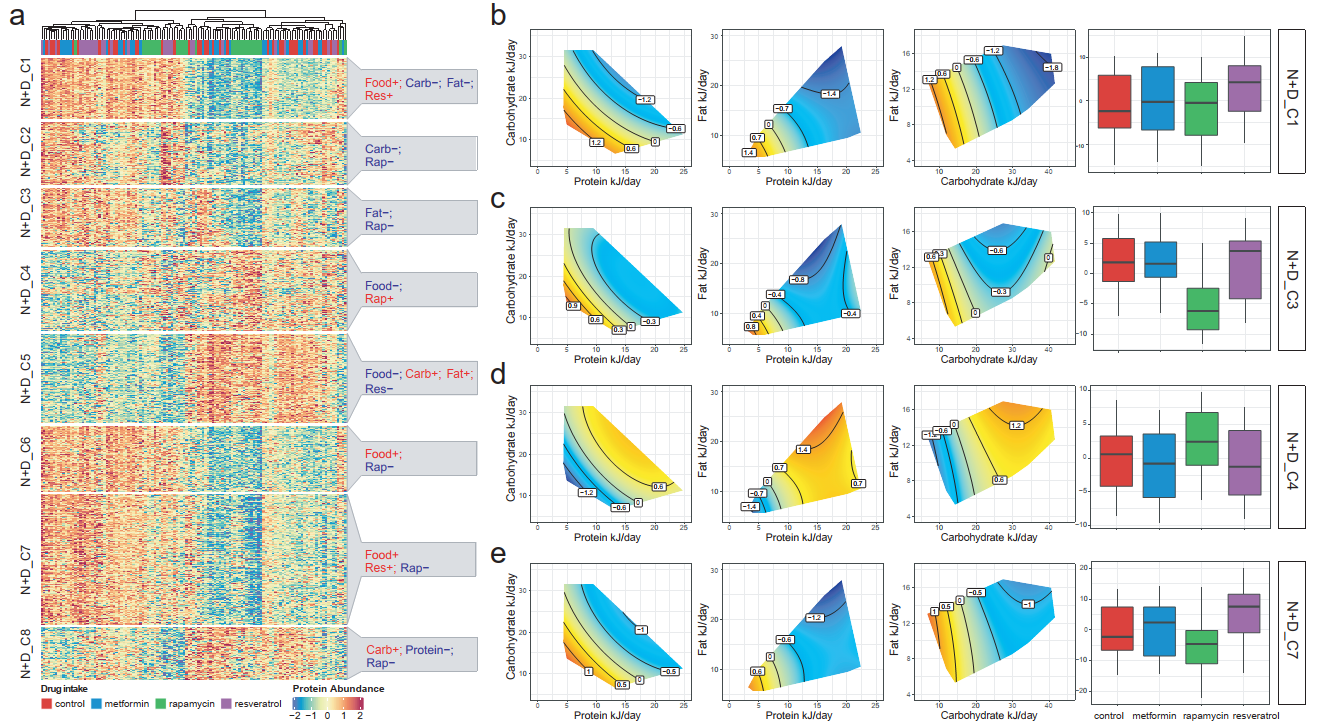

Fig S7. **Subclusters in "N+D" group and its annotation.** **a** The left panel shows the heatmap of protein abundance of proteins in the "N+D" group, split by subclusters. The right panel shows the annotation of each subcluster. **b** GFN and boxplot of PC1 of subcluster "N+D_C1". **c** GFN and boxplot of PC1 of subcluster "N+D_C3". **d** GFN and boxplot of PC1 of subcluster "N+D_C4". **e** GFN and boxplot of PC1 of subcluster "N+D_C7".

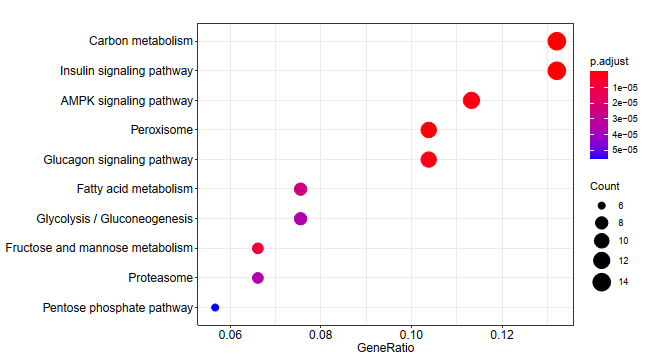

Fig S8. **Pathway enrichment analysis of subcluster "NxD_C4"**. Biologically meaningful pathways such as Insulin signaling pathway and AMPK signaling pathway are enriched.

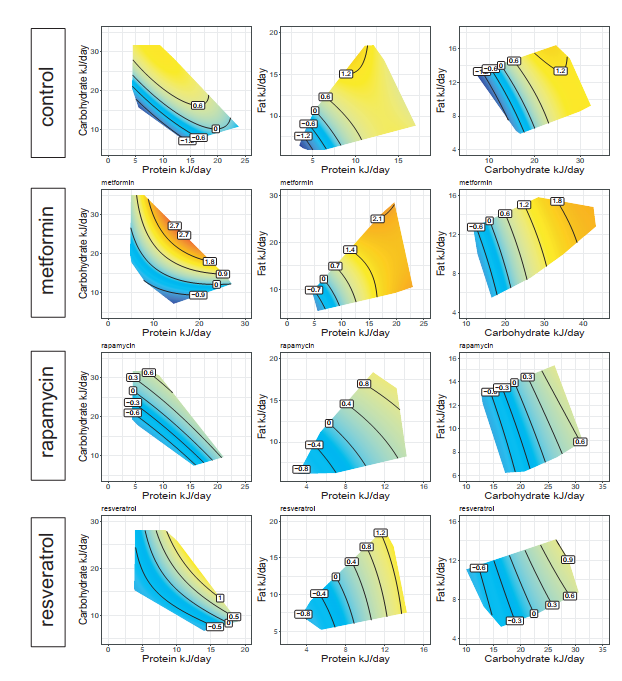

Fig S9. **GFNs of hub protein Pex11.** Xy-axis represents different combinations of nutrients and the contour shows the abundance of this protein. Each row corresponds to one type of treatment.

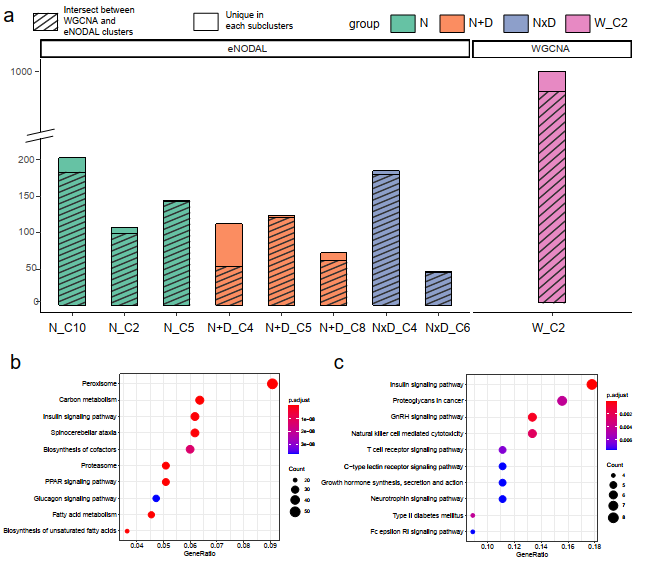

Fig S10. **Comparison between WGCNA clusters and eNODAL subclusters.** **a** Relationship between WGCNA cluster "W_C2" and related eNODAL subclusters. eNODAL panel shows a barplot of eight majority subclusters related to "W_C2", shaded bars represent that the proteins in each subcluster intersect with "W_C2". The WGCNA panel shows a bar plot of proteins in "W_C2"; the shaded bars represent that the proteins in "W_C2" intersect with all eight subclusters in eNODAL. **b** KEGG pathway enrichment analysis of subcluster "W_C2". **c** KEGG pathway enrichment analysis of subcluster "N+D_C8"
